# Biologically indeterminate yet ordered promiscuous gene expression in single medullary thymic epithelial cells

**DOI:** 10.1101/554899

**Authors:** F. Dhalla, J. Baran-Gale, S. Maio, L. Chappell, G. Holländer, C.P. Ponting

## Abstract

During thymic negative selection, medullary thymic epithelial cells (mTEC) collectively express most protein coding genes, a process termed promiscuous gene expression (PGE). Although PGE is crucial for inducing central T-cell tolerance, this process has not been established definitively as being either stochastic or coordinated. To resolve this question, we sequenced the transcriptomes of 6,894 single mTEC, including 1,795 rare cells expressing either of two tissue-restricted antigens, TSPAN8 or GP2. Transcriptional heterogeneity allowed partitioning of mTEC into 15 robustly-defined subpopulations representing distinct maturational stages and subtypes. Although 50 gene co-expression groups were robustly identified, few could be explained by chromosomal location, biological pathway, or tissue specificity. Further, GP2+ mTEC were randomly dispersed spatially within medullary islands. Thus although PGE exhibits ordered co-expression, biologically it is indeterminate. This likely enhances the presentation of diverse antigens to passing thymocytes during their medullary residency, while simultaneously maintaining mTEC identity throughout PGE.

## Introduction

Types of differentiated cells are distinguished by their restricted expression of transcription factors, upstream regulator proteins, and downstream target genes. If recapitulated out of context in other cell types, transcriptional programmes can induce the reprogramming of one mature somatic cell type into another (i.e. transdifferentiation) or trigger oncogenesis(Todd & Wong, 1999). Thymic epithelial cells (TEC), the major stromal cell constituent of the thymus(Barthlott *et al*, 2006; Takahama, 2006; Abramson & Anderson, 2017), express almost the entire protein coding genome(Sansom *et al*, 2014; Brennecke *et al*, 2015) and thus harbour an increased risk of transdifferentiation and consequently losing cellular identity. This capacity includes the competence to transcribe tissue-restricted genes (TRGs) whose expression in the periphery is normally limited to a single or small subset of tissues and genes whose expression is temporally or developmentally controlled or is sex-specific(Derbinski *et al*, 2001; Kyewski & Klein, 2006). This exhaustive transcriptional programme, termed promiscuous gene expression (PGE), provides a molecular mirror of the body’s self-antigens within TEC for the purposes of central T-cell tolerance induction.

T-cells that are unable to discriminate correctly between self- and non-self proteins risk provoking autoimmune disease. Therefore, during their intrathymic development, T-cells are subjected to stringent selection processes mediated by recognition of self-peptide::MHC complexes presented on the cell surface of TEC. TEC can be broadly categorized into cortical (c-) and medullary (m-) lineages based on their structure, anatomical location, molecular characteristics and functions(Rodewald, 2008; Vaidya *et al*, 2016). Studies examining TEC development and diversity have further revealed considerable heterogeneity within the mTEC compartment reflecting both maturational stages and functionally distinct mTEC subpopulations(Nishikawa *et al*, 2010; Metzger *et al*, 2013; Miragaia *et al*, 2018; Bornstein *et al*, 2018). During intrathymic selection, cTEC positively select thymocytes that express T-cell receptors (TCRs) capable of recognising peptide:MHC complexes. Following positive selection, cTEC and then mTEC remove potentially autoreactive T-cells bearing high-affinity TCRs for self-antigens via negative selection and mTEC additionally redirect those with intermediate affinity to a regulatory T-cell fate(Klein *et al*, 2009; Takahama, 2006; Sansom *et al*, 2014). Only 1-3% of thymocytes successfully fulfil the stringent criteria of thymic selection and exit to the periphery(Klein *et al*, 2014; Hogquist & Jameson, 2014).

PGE is partly under the control of the Autoimmune Regulator (AIRE), a transcriptional facilitator expressed in a subset of mature mTEC where it plays a role in the expression of just under 4000 genes(Sansom *et al*, 2014). Around 533 of these are entirely dependent on AIRE for their expression (AIRE-dependent), and the expression of the remaining 3,260 is enhanced in the presence of AIRE (AIRE-enhanced)(Sansom *et al*, 2014). AIRE-independent PGE further controls the expression of about 3,947 TRGs(Sansom *et al*, 2014).

Despite TEC expressing almost all protein coding genes at the population level, TRG expression at single-cell resolution is heterogeneous, with individual mature mTEC expressing only 1-3% of TRGs at one time(Sansom *et al*, 2014; Brennecke *et al*, 2015; Villaseñor *et al*, 2008; Derbinski *et al*, 2008; Meredith *et al*, 2015). A possible outcome of this mosaic expression pattern is the attainment of sufficiently high densities of particular self-antigen::MHC complexes on the cell surface of TEC to elicit tolerogenic signals within self-reactive thymocytes(Villaseñor *et al*, 2008).

Four molecular processes, not all mutually exclusive, could explain the heterogeneity of TRG expression within single mTEC. Type 1: TRG expression within single mTEC is entirely stochastic. Type 2: Different maturational stages or classes of mTEC activate TRG expression to different extents (with respect to breadth and/or level of gene expression) or, alternatively, activate different TRG subsets. Type 3: a programme of TRG co-expression otherwise evident in peripheral tissues is activated. In this scenario, TRGs whose expression is restricted to a particular tissue (e.g. liver) would be transcribed concurrently as a result of a transcriptional activation programme from that peripheral tissue being co-opted. This mechanism would arguably be the most hazardous with regards to the risk of transdifferentiation. As a potential example of the latter type, the recently identified tuft cell-like mTEC express a programme of genes contributing to the canonical taste transduction pathway(Bornstein *et al*, 2018; Miller *et al*, 2018). Type 4: TRGs are expressed co-ordinately owing to their physical co-location by one of two mechanisms: the loci are (a) positioned contiguously on the same chromosome or (b) distantly located but positioned adjacent via chromatin looping.

To resolve which of these four mechanisms contribute to the thymic representation of self-antigens we undertook large-scale single-cell RNA-sequencing of mTEC. Understanding how PGE is regulated within single TEC is essential for understanding the mechanisms by which these cells achieve their uniquely broad transcriptional programme without subverting their cellular identity. Previous studies have been limited by low cell numbers, with only several hundred mTEC analysed. We, therefore, undertook a study of the transcriptomes of thousands of single mTEC intending to resolve the existence and degree of PGE co-expression within them. Our selection of mTEC was both broad and narrow. The broad range of mTEC were unselected with respect to tissue-restricted antigen (TRA) expression; the narrow range contained two sets of mTEC that are rare in expressing TSPAN8 or GP2, two AIRE-regulated TRAs(Rattay *et al*, 2016; Sansom *et al*, 2014).

## Results

### Large scale single-cell RNA-sequencing data from FACS sorted mTEC

We chose to analyse the transcriptomes of single mTEC that were unselected or that promiscuously expressed either Tetraspanin 8 (TSPAN8) or Glycoprotein2 (GP2)(Rattay *et al*, 2016; Sansom *et al*, 2014) with the aim of adding statistical power to co-expression analyses. TSPAN8 is expressed in the gastrointestinal tract and several carcinomas(Agaësse *et al*, 2017; Zhu *et al*, 2017; Zhao *et al*, 2018) and GP2 is expressed in the pancreas and gastrointestinal tract(Cogger *et al*, 2017; Ohno & Hase, 2010); loss of tolerance to GP2 is associated with Crohn’s disease and primary sclerosing cholangitis(Werner *et al*, 2013; Tornai *et al*, 2018). Expression of both TRAs is enhanced in mTEC by the presence of AIRE and can be detected on their cell surface by flow cytometry. TSPAN8+ mTEC constitute approximately 7% and GP2+ mTEC about 2% of total mTEC (Figure 1a and S1) and each continue to actively transcribe their respective genes (Figure 1b).

**Figure 1.**
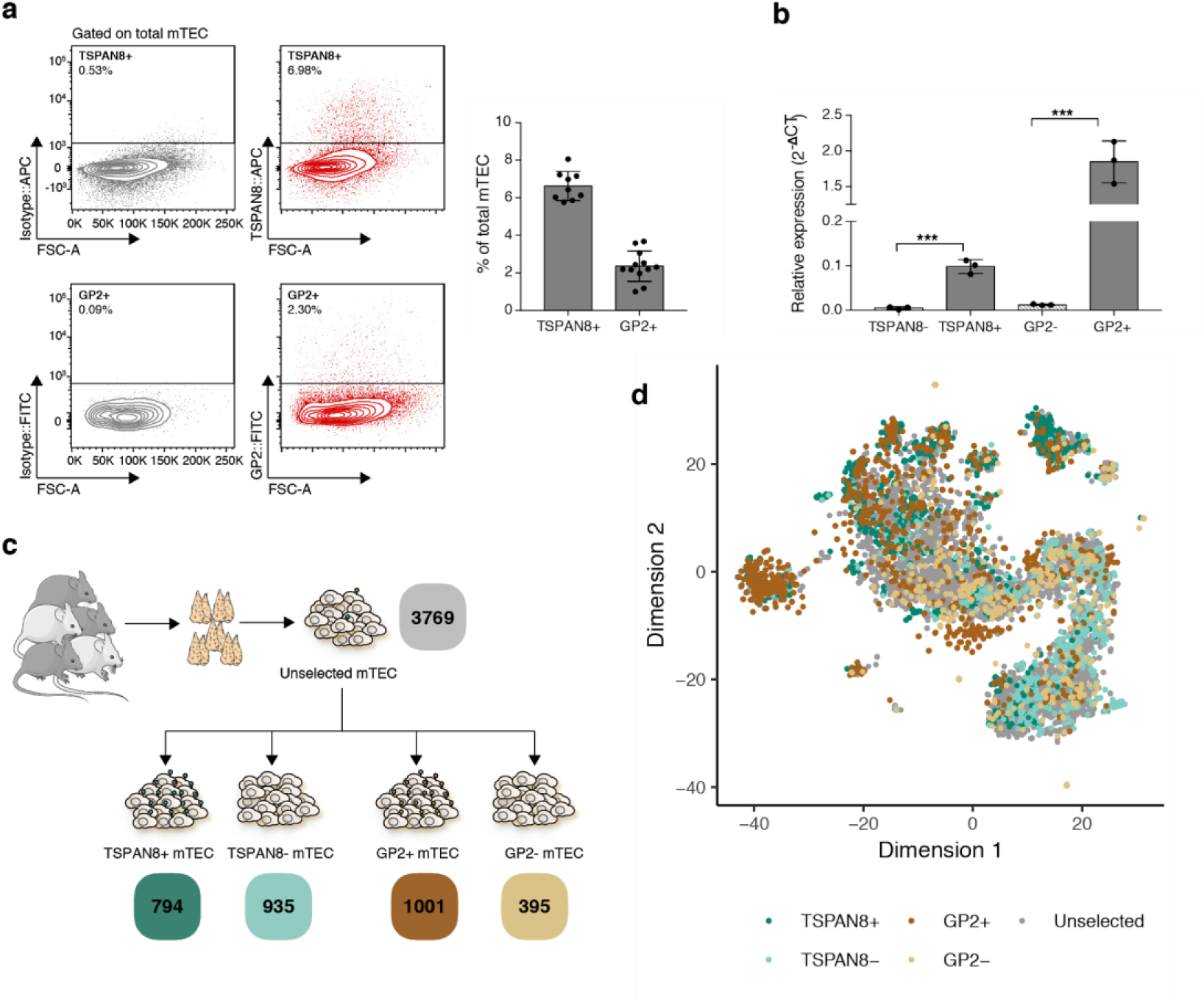
Deep transcriptome analysis at single-cell resolution of thousands of flow cytometrically sorted mTEC. (a) Left panels: mTEC promiscuously expressing TSPAN8 (top; TSPAN8+) or GP2 (bottom; GP2+) on their cell surface can be identified by flow cytometry. mTEC were identified as CD45-EpCAM+Ly51-(Fig. S1) and the gates for TSPAN8/GP2 (red) were set against an isotype control (grey). Right panel: Bar graph showing mean frequency (+/-sd) of TSPAN8+ or GP2+ cells within total mTEC; results represent pooled data from 3 (TSPAN8+) or 4 (GP2+) independent experiments each representing 3 individual mice. (b) Identification of TSPAN8 or GP2 protein expression via FACS reflects mRNA expression. Bar graph showing mean expression (+/-sd) of *Tspan8* and *Gp2* mRNA relative to *β-actin* by RT-qPCR on FACS sorted mTEC negative or positive for TSPAN8 or GP2 protein, respectively; n=3, representative of 2 independent experiments. (c) Schematic representation of cell populations sorted by flow cytometry for single-cell RNA-sequencing. Numbers of recovered cells are indicated below, coloured by category. (d) t-SNE visualisation of mTEC subpopulations from all experiments coloured by surface phenotype established via flow cytometry; see Figure 2B, C for the distribution of unselected mTEC.

mTEC sequencing libraries were derived from 15 individual female mice across three independent experiments. To examine strain-specific patterns in mTEC gene expression, C57BL/6 (n=9) and BALB/c (n=2) mice were investigated as well as their F1 cross, C57BL/6 x BALB/c (n=4).

The transcriptomes of 6,894 single mTEC were analysed including 794 TSPAN8+, 935 TSPAN8-, 1,001 GP2+, 395 GP2-, and 3,769 unselected mTEC (Figure 1c) making this the largest single-cell RNA-seq dataset investigating PGE in mTEC. Together, these cells showed expression of 22,819 genes encoded across all chromosomes (except Y, as expected), including 19,091/21,663 or 88% of protein coding genes. We further categorized these transcripts according to their dependence on AIRE (as defined in Sansom *et al*. 2014) and observed the expression of 89% of AIRE-dependent (N=477), 98% of AIRE-enhanced (N=3,210) and 94% of AIRE-independent TRGs (N=3,720; Figure S2a). Therefore, the single cells largely recapitulated expression observed in TEC population-level analyses(Sansom *et al*, 2014). A median of 1,830 genes was detected per cell of which 50 (median) were AIRE-regulated TRGs (Figure S2c and d).

### Cell subpopulations were robustly identified across independent datasets

The transcriptomes of the 6,894 mTEC were projected into a reduced dimensional space resulting in a large, contiguous, central ‘body’ of cells surrounded by several ‘satellite’ clusters (Figure 1d). Within the central body, the majority of mTEC fell along a manifold characterised by a transition from predominantly TSPAN8- or GP2-mTEC at the lower right pole, to TSPAN8+ or GP2+ mTEC at the upper left pole (Figure 1d, Figure 2). TSPAN8+ and GP2+ mTEC each contributed to distinct satellite clusters (Figure 2 green and brown arrows in panels a, b, d). This pattern indicates that subsets of mTEC expressing a particular TRA express distinct transcriptomes non-stochastically that differentiate them from most other mTEC, a finding that is inconsistent with a Type 1 mechanism (entirely stochastic TRG expression).

**Figure 2.**
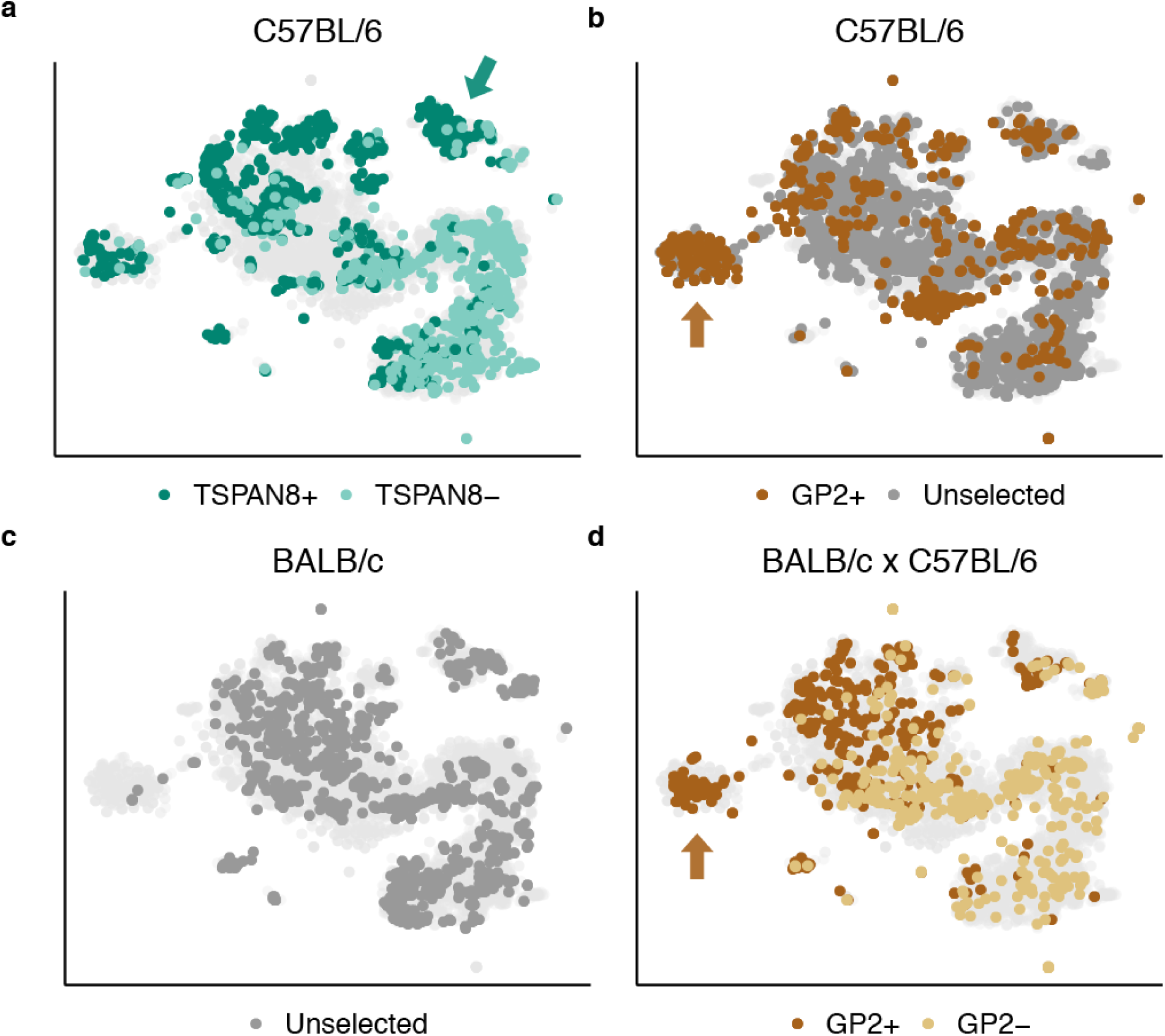
Cell subpopulations are reproducible across different experiments, mouse strains and cell surface phenotypes. Panels (a-d) are t-SNE visualisations of individual datasets from the current study, overlaid on the combined dataset (light grey). Each dot represents an individual mTEC coloured as follows: Unselected: dark grey; TSPAN8+: dark green; TSPAN8-: light green; GP2+: dark brown; GP2-: light brown. (a) 794 TSPAN8+ and 935 TSPAN8-mTEC from C57BL/6 mice; green arrow identifies TSPAN8 preferred cluster. (b) 549 GP2+ and 2561 unselected mTEC from C57BL/6 mice; brown arrow identifies GP2 preferred cluster. (c) 1208 unselected mTEC from BALB/c mice. (d) 452 GP2+ and 395 GP2-mTEC from BALB/c x C57BL/6 F1 mice; brown arrow identifies GP2 preferred cluster. The whole space has good representation in most samples with the notable exception of the GP2+ enriched region (brown arrow in panel B&D), which is underrepresented in the unselected cells from BALB/c due to a combination of fewer TEC analysed and the lack of enrichment for antigen-positive TEC (such as in the TSPAN8+ or GP2+ experiments).

Data were collected from three independent experiments probing different strains of mice and cell phenotypes (Table S1). Nevertheless, each of the conditions captured the transcriptomic diversity of the batch-corrected meta-experiment (Methods; Figure 2). The transition from TRA-to TRA+ mTEC surrounded by satellite clusters is additionally evident in our analysis of published single-cell TEC datasets (Figure S3)(Sansom *et al*, 2014; Brennecke *et al*, 2015; Miragaia *et al*, 2018; Bornstein *et al*, 2018). We conclude that a non-random and robustly defined set of diverse mTEC subpopulations is reproducible across multiple distinct experiments using different mouse strains and single-cell experimental protocols and that TRG expression biases are apparent across subpopulations.

### Heterogeneity within the main mTEC body reflects their cellular maturation trajectory

Next, we considered the cells’ maturational states and found that a Type 2 process (see Introduction) best explains PGE within single mTEC. Using unsupervised clustering, these cells resolved into 15 distinct subpopulations (Figure 3a) which were reproducible and robust across mouse strains and unselected or selected (TSPAN8+ or GP2+) mTEC (Figure S4b). These subpopulations were defined by genes whose expression varies throughout mTEC maturation (Figure 3). Cluster definitions were largely preserved when mTEC were clustered using all genes, or only TRGs, or when excluding TRGs (Figure S4c).

**Figure 3.**
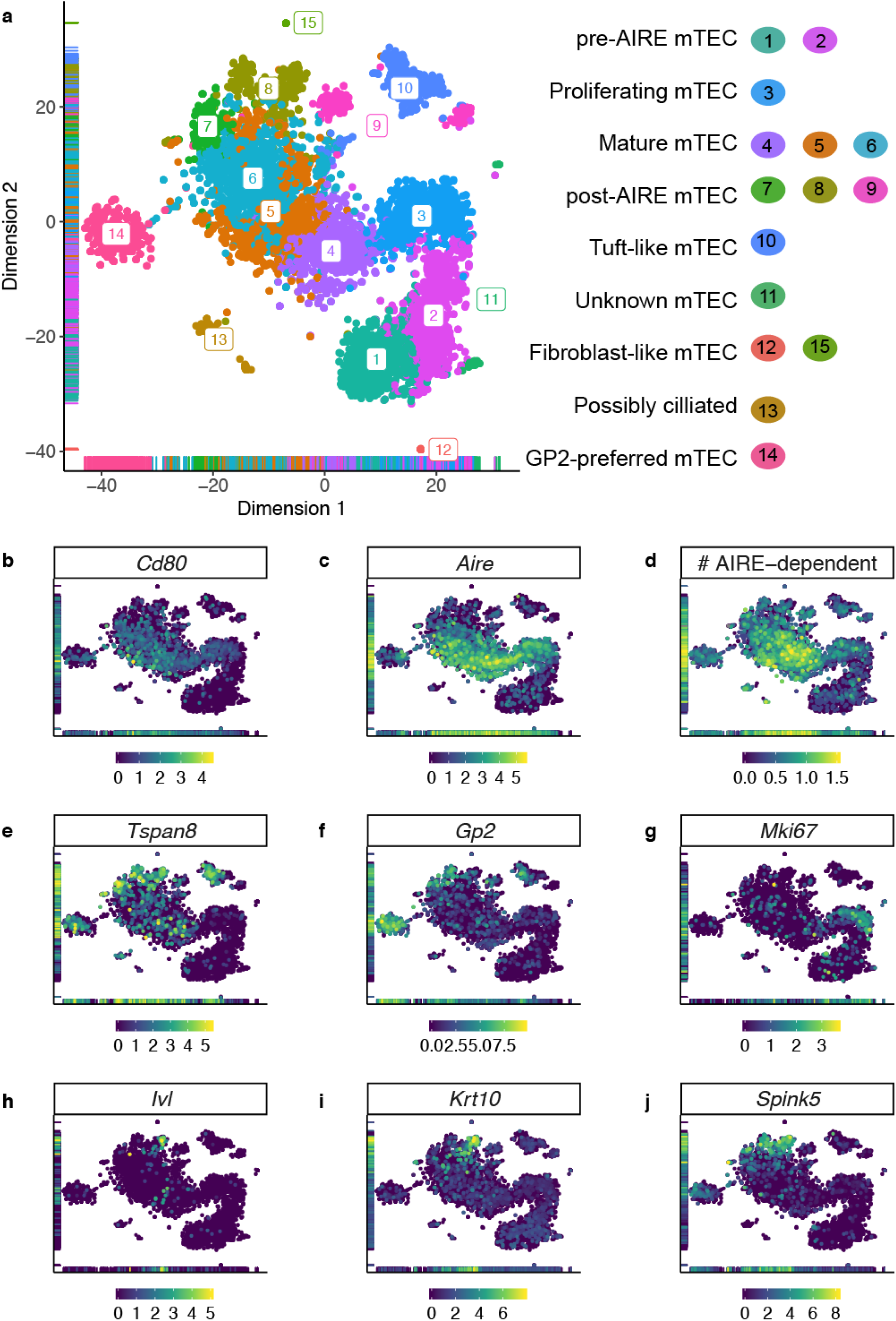
Cell subpopulation clusters recapitulate known features of mTEC maturation. (a) t-SNE visualisation of mTEC subpopulations. Each dot represents a cell coloured by cluster number. (b-c, e-j) Log2 expression level of *Cd80, Aire, Tspan8, Gp2, Mki67, Involucrin* (*Ivl*), *Keratin 10* (*Krt10*) and *Spink5* across the dataset. (d) Log10 of the number (#) of AIRE-dependent genes expressed per cell.

We begin with the mTEC of **clusters 1** and **2** (lower right of Figure 3a). These were mainly TSPAN8-, GP2-, or unselected mTEC (Figure 2 and S4a) and had little-to-no mRNA expression of *Aire* or AIRE-regulated TRGs, including *Tspan8* and *Gp2* (Figure 3c-f). Because these clusters expressed *Pdpn* and *Ccl21a* (Figure S5), they likely represent immature junctional(Onder *et al*, 2015) and pre-AIRE mTEC(Michel *et al*, 2017). **Cluster 3** also contained mostly TSPAN8-, GP2-, or unselected mTEC (Figure S4a) that expressed *Mki67* and *Aire* (Figure 3g and c). These were predicted(Scialdone *et al*, 2015) to be in the G_2_/M-phase of the cell cycle based on their gene expression profile. This cluster could, therefore, represent a proliferating subpopulation or maturational stage of mTEC. **Cluster 4** represents the next likely maturational stage as these mTEC (i) were mostly negative for expression of TSPAN8 or GP2 at both protein and mRNA level and (ii) highly expressed *Aire* (Figure 3c, e, f, S4a) and hence also transcripts for AIRE-regulated TRGs (mean of 90 per cell). **Clusters 5** and **6** contained mTEC with the broadest TRG representation: collectively they expressed approximately 98% of detected TRGs. These mTEC not only expressed *Aire* (Figure 3c) and a high number of AIRE-regulated TRGs (mean of 82 per cell in cluster 5 and 72 per cell in cluster 6), but also *Cd80* (Figure 3b) and *Cd86* (Figure S5). Moreover, they expressed TSPAN8 or GP2 protein and mRNA more frequently than clusters 1-4 (Figure 3e, f; Figure S4a). These features identified clusters 5 and 6 as typical representatives of PGE competent mTEC(Derbinski *et al*, 2005). The mTEC in **clusters 7** and **8** also expressed TSPAN8 and GP2 protein and mRNA (Figure 3e, f; S4a) and a moderate number of AIRE-dependent TRGs (Figure 3d), but they had reduced or no expression of *Aire, Cd80* and *Cd86* (Figure 3b-c and S5). In addition, they expressed markers associated with epithelial cell terminal differentiation including *Ivl* and *Krt10(Michel et al, 2017)* (Figure 3h-i), and *Spink5* (Figure 3j). The latter has previously been found in Hassall’s corpuscles(Galliano *et al*, 2005; Bitoun *et al*, 2003) and appears to be a more informative indicator of terminally differentiated mTEC in our dataset than the classically used *Ivl* and *Krt10.* In keeping with a terminally differentiated phenotype, TSPAN8 or GP2 protein positive mTEC were also significantly enriched for DSG3 expression (Figure S6), another marker associated with epithelial cell terminal differentiation found in Hassall’s corpuscles(Wang *et al*, 2012; Wada *et al*, 2011).

Consistent with cells of the main body of mTEC transitioning from TSPAN8-/ GP2-at its lower right pole to TSPAN8+ / GP2+ at its upper left pole, these cells also line up along a trajectory in diffusion space (Figure 4a). We ordered the cells in pseudotime using this trajectory. In this first analysis, the two main branches in the trajectory originated at **cluster 3**, which we have inferred to be proliferating mTEC based on their expression of cell cycle relevant transcripts including *Mki67* (Figure 3g). From cluster 3, mTEC were predicted to proceed either to **cluster 4** and then **cluster 7** (Figure 4b) or **8** (Figure 4c), or alternatively, progress via **cluster 2** to **cluster 1** (Figure 4d). An orthogonal method that uses pre- and post-spliced mRNA reads to order cells(La Manno *et al*, 2018), produced a concordant set of trajectories, with one exception, namely that the proliferating mTEC in **cluster 3** appeared to derive from **cluster 2** (Figure 4e). Taken together, these results suggest that proliferating mTEC in **cluster 3** and *Aire*+ mTEC in **clusters 4-6** originated from the *Aire-Cd80-CD86-* mTEC in **cluster 2**. The *Aire*-*Cd80-Cd86-* mTEC from **clusters 7-8** appeared to derive from mature mTEC of **clusters 5-6**(Yano *et al*, 2008; Michel *et al*, 2017; Wang *et al*, 2012) and were transcriptionally distinct from the *Aire-Cd80-Cd86-* cells in **clusters 1** and **2**. Consequently, we propose that **clusters 1** and **2** represent pre-AIRE mTEC (distinguished by *Ccl21a* and *Pdpn* expression) while **clusters 7** and **8** represent post-AIRE mTEC (distinguished by *Ivl, K10* and *Spink5* expression). These findings are in keeping with current models of mTEC maturation(Sun *et al*, 2013; Michel *et al*, 2017)

**Figure 4.**
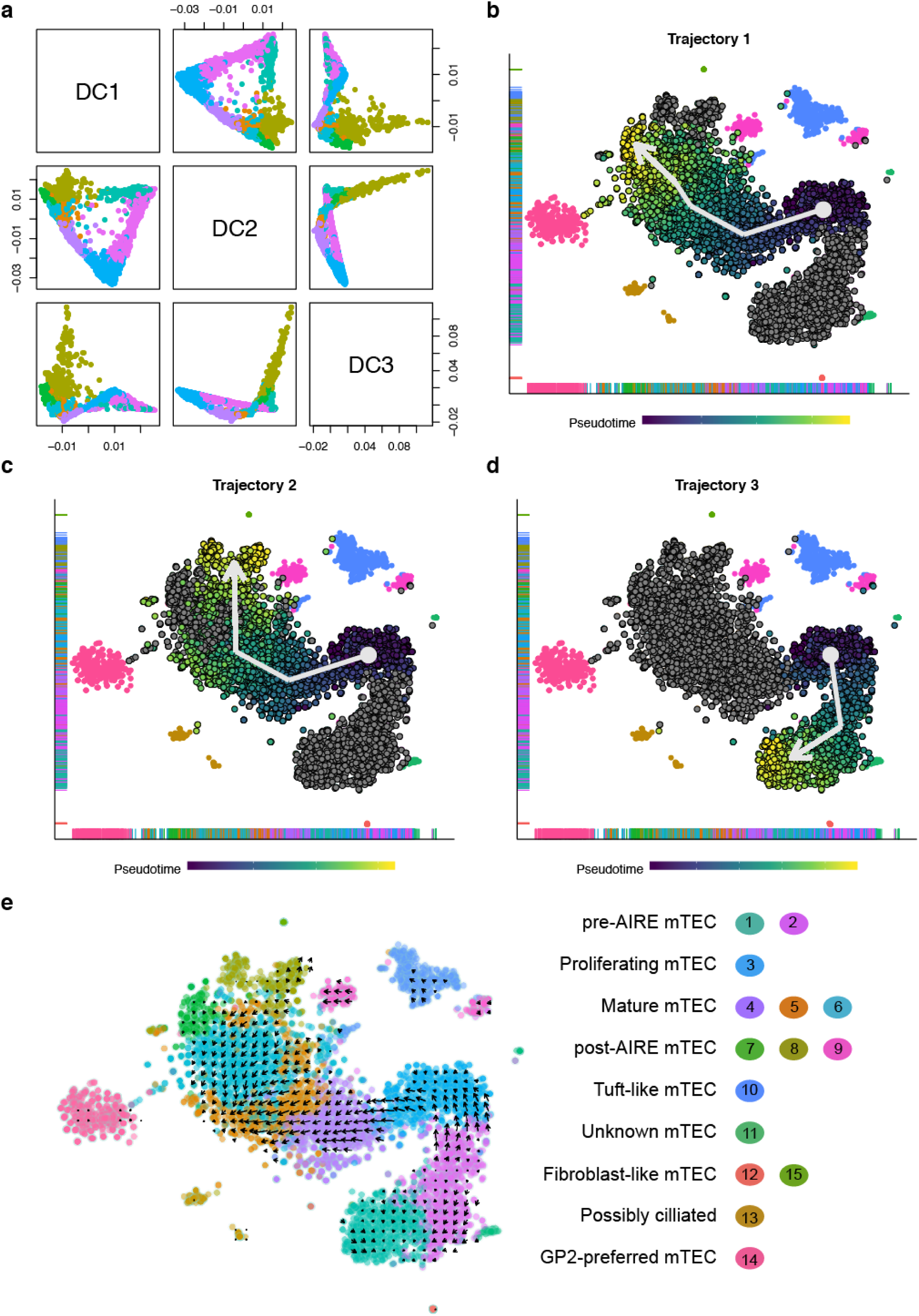
Pseudotemporal ordering of mTEC resolves three trajectories of mTEC maturation. (a) Paris plot of the first three diffusion components (DC) of the dataset. (b-d) t-SNE visualisation of mTEC subpopulations. Each dot represents a cell coloured by the inferred position in the pseudotemporal ordering. A white arrow indicates the direction of each trajectory. (e) t-SNE visualisation of mTEC subpopulations with arrows showing the ordering as identified by RNA velocity analysis. Each dot represents a cell coloured by cluster ID (right).

Two of our main observations are, to our knowledge, novel. Firstly, pre-AIRE mTEC can be distinguished into two subtypes (clusters 1 and 2), and secondly the mTEC in **cluster 2** precede those in **cluster 1** in the pseudotime analysis. One potential interpretation of this is that the **cluster 1** mTEC represent a class of quiescent progenitor TEC(Wong *et al*, 2014), while **cluster 2** mTEC represent MHCII^lo^ mTEC that are transitioning to MHCII^hi^ mTEC. The mTEC of **cluster 1** highly express genes such as *Itga6* (CD49f) and *Ly6a* (Sca-1) that are markers of a quiescent mTEC progenitor population with limited regeneration potential(Wong *et al*, 2014).

### Aire-/low satellite clusters lie separately from the main trajectory of mTEC maturation

Seven satellite clusters surround the main body of mTEC as displayed by the tSNE visualisation (Figure 3a). **Cluster 9** (split into 2 sub-clusters) appeared to contain mTEC involved in negative regulation of proliferation (Gene Ontology analysis) and in pathways dealing with the response to stress (Reactome analysis; Figure S7). These mTEC were a mixture of TSPAN8 and GP2 positive and negative cells and were largely devoid of *Aire* expression. **Cluster 10** contained TEC recently labelled as “thymic tuft cells”(Bornstein *et al*, 2018; Miller *et al*, 2018). This cluster was highly populated by TSPAN8+ mTEC (Figure 2a and S4a) and was characterised by the expression of *Pou2f3*, a transcription factor involved in regulating tuft cell function in the intestinal epithelium and respiratory tract (Figure S5)(Yamashita *et al*, 2017; Reid *et al*, 2005) and its target genes including *Tas2r* genes and *Trpm5(Yamashita et al, 2017)*. **Cluster 11** was characterised by the expression of genes related to RNA metabolism and nonsense-mediated decay (Figure S7) and **cluster 12** by organisation of the extracellular matrix (Figure S7). Both clusters 11 and 12 were mainly TSPAN8- and GP2- mTEC. **Cluster 13** cells express genes involved in the response to stress and external stimuli as well as cilium assembly (Figure S7) and are mostly TSPAN8+ or GP2+ mTEC. **Cluster 14** contained an over-representation of GP2+ mTEC and expressed low levels of *Aire*. Other markers of this cluster included the chemokine ligands *Ccl6* (Figure S5), *Ccl9* and *Ccl20* and chemokine receptor type 5 (*Ccr5)* suggesting a potential role in cell communication. Finally, **Cluster 15** was characterised by the expression of genes related to the organisation of the extracellular matrix (Figure S7).

Using FACS to enrich for TSPAN8+ mTEC and GP2+ mTEC, respectively, ensured that we investigated a large number of rare **cluster 10** and **14** cells. Nearly half the mTEC in these clusters were positive for their respective TRAs (44% and 49%, respectively). Importantly, these clusters were robust to clustering of unselected mTEC alone (Figure 2c). Furthermore, while **cluster 10** contained thymic tuft cells(Bornstein *et al*, 2018; Miller *et al*, 2018), **cluster 14** was transcriptionally distinct and expressed a set of chemokine ligands and receptors that are absent from **cluster 10**.

These observations argue for an uneven expression of TRGs across mTEC subpopulations, implying that satellite clusters show preference for expression of particular gene subsets and providing additional evidence against a Type 1 process (TRG expression is stochastic), and in favour of a Type 2 process (different maturational stages or classes of mTEC activate TRG expression differentially).

### Gene module clustering reveals robustly identified gene co-expression groups supportive of PGE being an ordered process

The robust identification of 15 distinct mTEC clusters suggested an ordered process that selects TRGs to be co-expressed within single mTEC. To further investigate the patterns of gene co-expression, we applied a method of gene module clustering that accentuates expression similarities found among rarely observed genes and attenuates similarities found among frequently observed genes. This allowed us to assign genes to single gene modules that exhibited a distinctive gene co-expression profile across our dataset. In total 14,861 genes were assigned to 50 modules (Figure 5a). A further 7,958 genes, including many olfactory and vomeronasal receptor genes, could not be assigned to a module due to their low detection frequency. The definition of these 50 gene modules was both reproducible and robust because re-clustering of 100 random subsamples of the data produced highly similar modules (mean adjusted mutual information (AMI) score = 0.695; Figure 5b). This finding is not explained by cellular expression levels because the analysis used a TF-IDF transform(Manning *et al*, 2008) ensuring that frequently expressed genes do not contribute substantially to the module identities.

**Figure 5.**
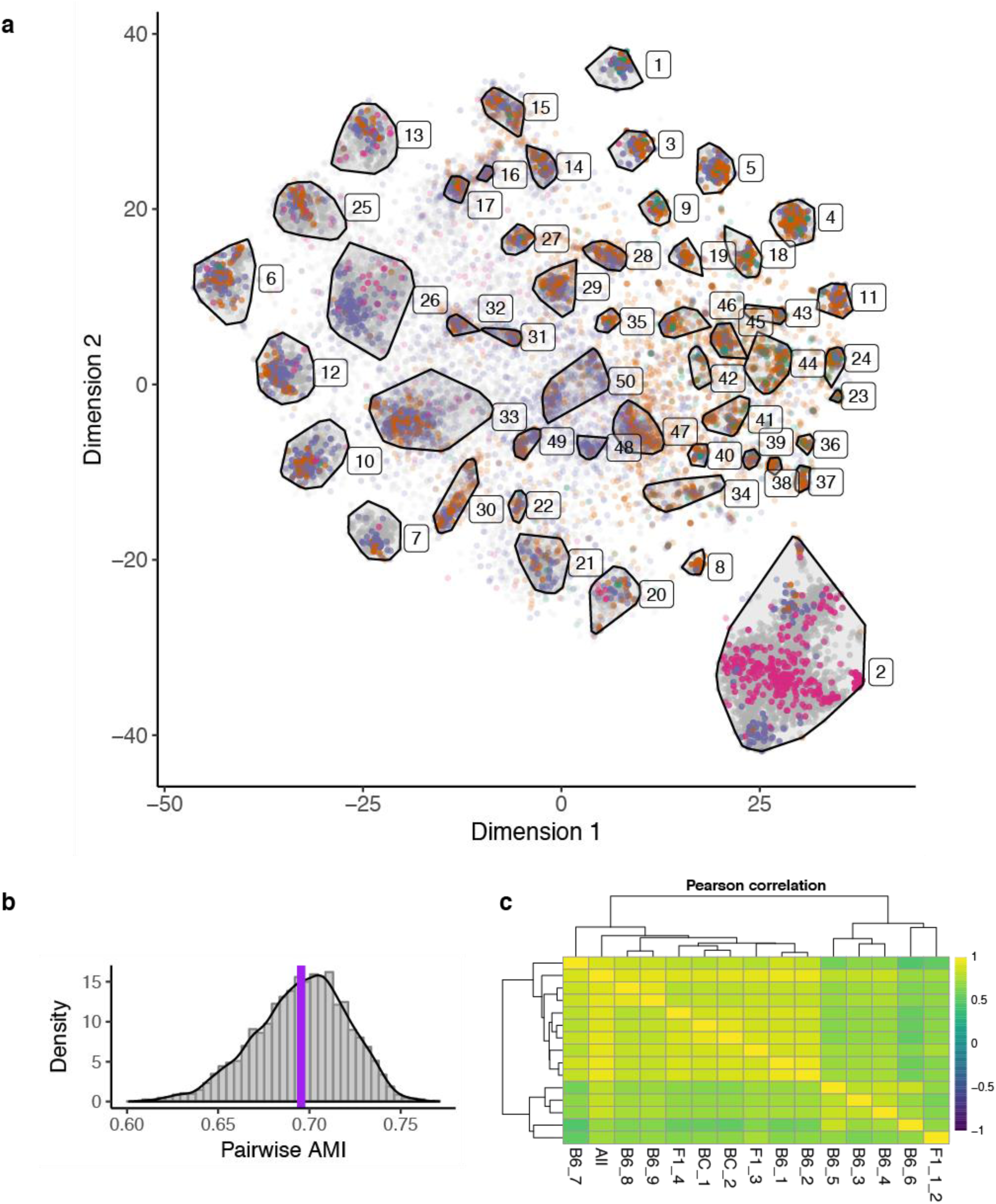
Robust gene co-expression modules indicate that mTEC gene co-expression has order. (a) t-SNE visualisation of the partition of 22,819 genes into 50 mutually exclusive gene modules. Each dot represents a gene and is coloured according to gene expression category (AIRE-dependent, AIRE-enhanced, AIRE-independent TRG, Other, Housekeeping, Unclassified). The colour intensity of each dot/gene is proportional to its membership probability within each module. (b) Histogram of adjusted mutual information (AMI) between random samples. The mean AMI value is indicated by a vertical purple line; AMI values lie between 0 and 1. (c) Pearson correlation of co-expression frequency for pairs of TRGs (AIRE-regulated and AIRE-independent) between individual mice or all mice pooled together (see Table S1 for mouse identifiers). Mean correlation = 0.77, with the largest divergence observed between samples with higher read depth (B6_3-B6_6, and F1_1_2).

Next, we sought to determine the variability of TRG co-expression between individual mice (of either the same or different genetic backgrounds). For each pair of TRGs, the fraction of mTEC expressing both TRGs was calculated per mouse, these fractions were then compared across all mice in our dataset. The mean Pearson correlation of these fractions across all mouse pairs was high (average value of 0.77; Figure 5c). Our findings thus demonstrate that sets of TRGs were repeatedly co-expressed in individual mTEC and that this TRG co-expression was replicated both in random subsets of mTEC and across different mice.

Half of the 50 gene modules were significantly enriched in TRGs (53% median proportion of AIRE-regulated or AIRE-independent TRGs across these modules; Figure 6a-c). By contrast, 7 gene modules had significantly fewer TRGs than expected by chance (15% median TRG proportion). Genes in each module were expressed at a comparable level (normalised UMI count) and within similar mTEC subsets across the 6,894 single cells analysed. These observations imply that complex gene co-expression programmes were replicated in individual modules across many mTEC and were driven by both TRGs and by non-TRGs. The significant contribution of TRGs to half of the gene modules further implied that their co-expression substantially defined these modules. For example, *Gp2* was assigned to module 7 and genes in module 7 were most highly expressed in the post-*Aire* clusters 7 and 8, as well as in the GP2-preferred cluster 14. In contrast, *Tspan8* was assigned to module 31, whose member genes were most highly expressed in the tuft cell-like cluster 10. Consequently, although both TSPAN8+ and GP2+ mTEC were located in some of the same cell clusters, *Tspan8* and *Gp2* were co-expressed with very different sets of TRGs. This further supports the theory that PGE results in ordered gene co-expression.

**Figure 6.**
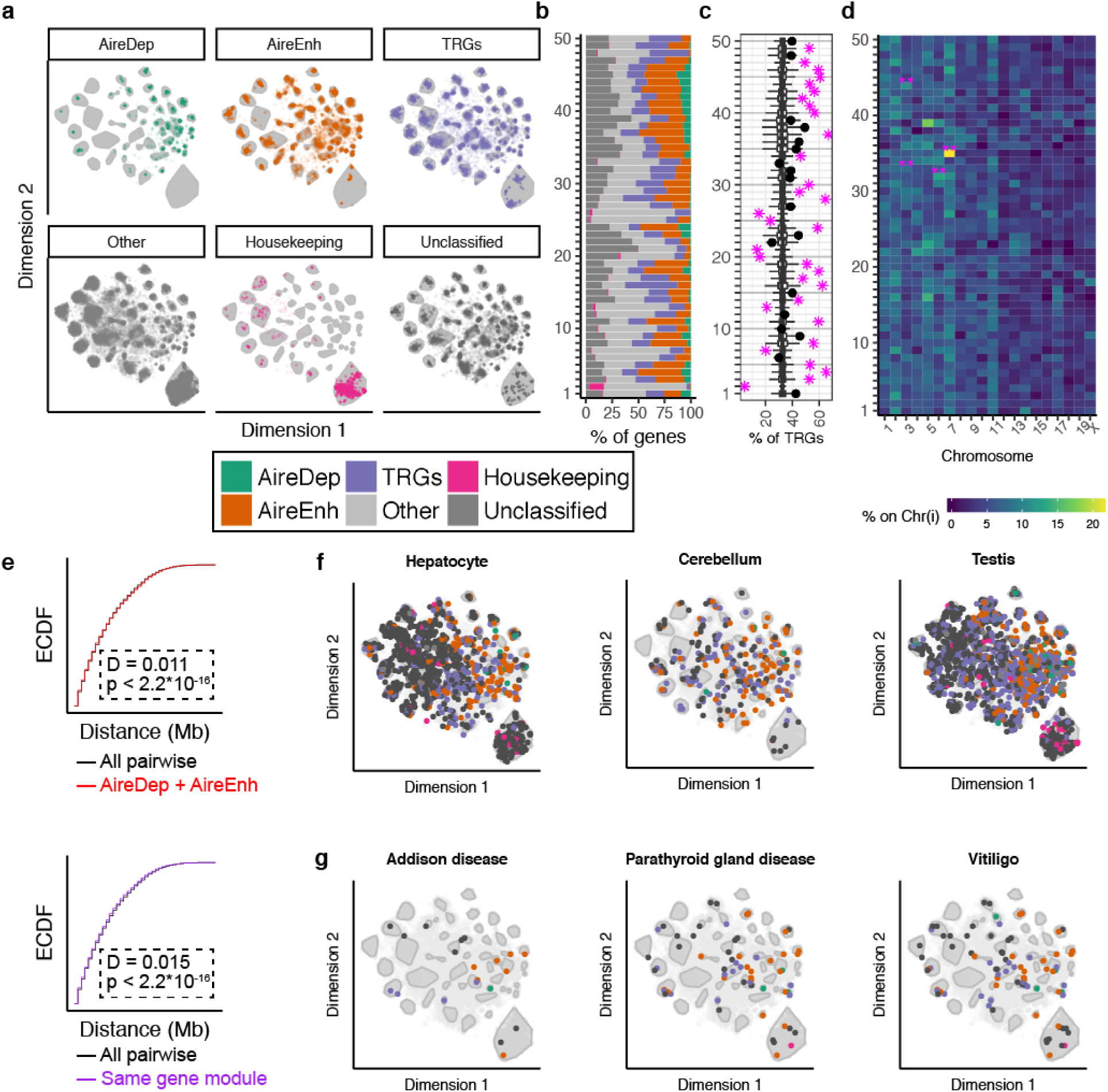
Randomness underlies the structure in gene co-expression modules. (a) Visualisation of the composition of gene modules coloured by gene expression category (AIRE-dependent, AIRE-enhanced, AIRE-independent TRG, Other, Housekeeping, Unclassified). (b) Gene expression category composition of each gene module displayed as a circular stacked bar-chart. (c) Percentage of TRGs in each module. Modules significantly enriched or depleted in TRGs are indicated by a magenta star and non-significant modules by a black dot. Boxplots show the expected contribution of TRGs from 10,000 random samples. (d) Chromosomal locations for genes expressed within each gene module. ** Adjusted p-value < 0.01 (in pink) indicates that more genes assigned to a given module (x-axis) are located on a particular chromosome (y-axis) than expected by chance (empirical p-value, corrected for multiple comparisons). (e) Empirical cumulative distribution function (ECDF) for distribution of the pairwise genomic distance between co-expressed genes. Top: all pairwise distances in grey; between AIRE-regulated genes only in red; Bottom: all pairwise distances in grey; between genes within the same gene module only in purple. (f) Distribution of tissue-specific genes across gene modules for hepatocytes (left), cerebellum (middle) and testis (right); tissue-specific genes are highlighted with a coloured dot that denotes the gene category. (g) Distribution of disease-related auto-antigen genes across gene modules for parathyroid gland disease (left), Addison’s disease (middle) and vitiligo (right); auto-antigen genes are highlighted with a coloured dot that denotes the gene category.

Three modules contained genes frequently expressed in mTEC that were not significantly enriched for TRGs (Figure 6b-c). One of these modules (module 2), contained genes that were detected, on average, in approximately 45% of mTEC, whereas genes in modules 26 and 48 were expressed in about 12% of mTEC and all three contained few AIRE-regulated TRGs. Several modules contained genes highly expressed in a particular mTEC maturational state or subpopulation. For example, module 26 contained genes expressed in the cycling cells of cluster 3 (including *Mki67*, cyclins and *E2f* genes) and module 48 encompassed genes expressed in the *Aire-*expressing clusters 4 and 5. Modules 31 and 32 included genes that are characteristic of thymic tuft cells (cluster 10) such as the *Tas2r* family, *Il10, Il25* and *Dclk1*, and genes co-expressed under the transcriptional control of POU2F3(Yamashita *et al*, 2017)(Figure S5). Module 49 contained transcripts of chemokines (including *Ccl6, Ccl9*, and *Ccl20*) and chemokine receptors (*Ccr1, Ccr2* and *Ccr5)* typical of cluster 14 (Figure S5).

In summary, 50 gene co-expression modules were identified, half of which were largely driven by TRG co-expression patterns reproducible in different mice. This again argues against a stochastic mechanism for TRG expression within single mTEC (Introduction).

### Whilst gene expression in mTEC is ordered, *gene membership in co-expression clusters is biologically indeterminate*

Next, we asked what feature, such as chromosomal location, intergenic distance, tissue-, pathology- or pathway-restricted expression, might explain the observed order of TRG co-expression in single mTEC. The co-expression pattern identified in the majority of the gene modules was nearly always independent of expression by a single chromosome (Figure 6d), with the exception of modules 33 and 44, which contained more transcripts of genes located on chromosome 3, and modules 32 and 35, which had a higher frequency of transcripts from genes on chromosomes 6 and 7 than expected. Nevertheless, such enrichments are not highly explanatory of the co-expression order observed in 46 of the 50 gene modules.

The chromosomal distances between genes present within the same gene module were near identical to those of all other genes regardless of AIRE-dependency (Figure 6e). Moreover, we found no evidence that individual modules were biased in their gene expression for (i) TRAs of individual peripheral tissues (Figure 6f), (ii) antigens characteristic of individual organ-specific autoimmune pathologies (Figure 6g), or (iii) molecules assigned to a particular cellular pathway or Gene Ontology term. These findings are not consistent with a Type 3 mechanism in which mTEC recapitulate the transcriptional programme of a peripheral tissue.

Finally, we considered whether the transcriptomic identity of mTEC varied according to their spatial location. For this we computed the G-function, the cumulative distribution function of distances between nearest neighbour mTEC each expressing the same TRG, here GP2 (Figure 7a-b). For comparison, G-functions were also computed for simulated mTEC that are: (i) evenly spaced (ii) dispersed randomly, or (iii) clustered (Figure 7c). Our results showed that GP2+ mTEC are spatially distributed within the medulla in a manner consistent with a random process (Figure 7a-b and Figure S8).

**Figure 7.**
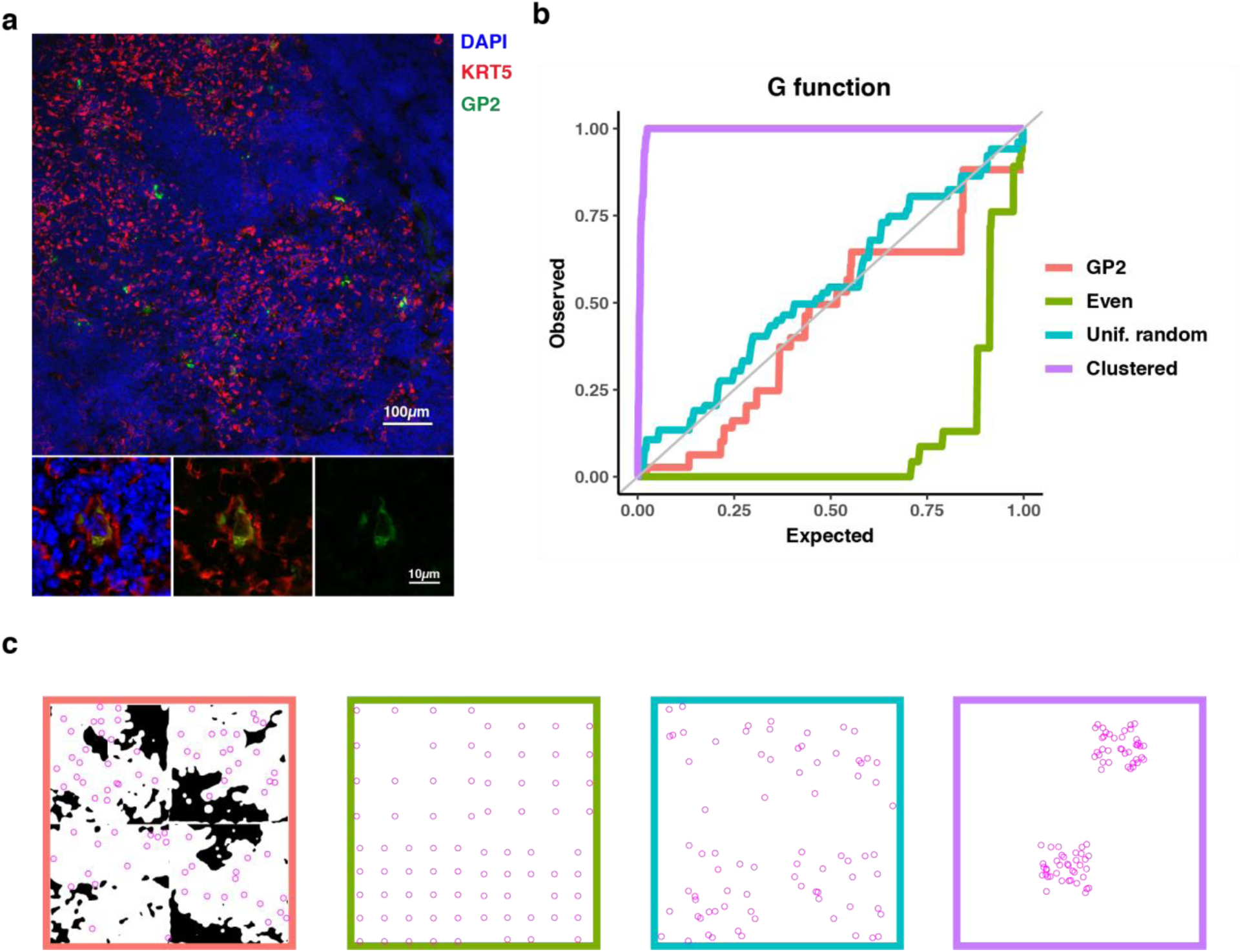
GP2+ mTEC are not spatially clustered but instead dispersed randomly within medullary islands. (a) Representative microscope image of a thymic slice. GP2 is stained in green and KRT5 (mTEC) in red. Zoomed images show the overlap of GP2 and KRT5 for one Gp2+ mTEC. (b) Observed versus expected G-function for GP2 spacing (red) and exemplar spacings. (c) Masked medullary region (white) for four slides stained as in (a) with GP2+ mTEC as magenta circles. Exemplar distributions for comparison (coloured as in b).

Together these results indicate that PGE in the medulla is a biologically indeterminate yet ordered process whose order is provided by repeated co-expression of particular gene subsets. Furthermore, these co-expressed genes are not systematically collocated in the linear genome and are not linked by tissue specificity, or by biological or disease processes.

## Discussion

The molecular processes underlying PGE in single mTEC have remained unclear despite numerous studies addressing this topic. These studies, which have been limited by technology and cell number, have not established conclusively whether TRG expression in single mTEC is stochastic or ordered(Sansom *et al*, 2014; Derbinski *et al*, 2005, 2008; Villaseñor *et al*, 2008; Pinto *et al*, 2013; Brennecke *et al*, 2015; Rattay *et al*, 2016; Meredith *et al*, 2015). Some studies that failed to detect co-expression patterns used single-cell PCR limited to the detection of only a very small number of TRGs in a small number of cells (Derbinski *et al*, 2008; Villaseñor *et al*, 2008). Co-expression patterns were detectable in three studies that either preselected mTEC for the expression of cell surface-expressed TRAs(Pinto *et al*, 2013) or conducted single-cell RNA-sequencing of hundreds of single mTEC (Brennecke *et al*, 2015; Meredith *et al*, 2015). Two studies found that mTEC express a larger repertoire of TRGs as they mature and that single mTEC are not restricted to expressing TRGs belonging to particular peripheral tissues, biological pathways or controlled by the same transcription factors (Pinto *et al*, 2013; Meredith *et al*, 2015).

Our analysis of thousands of single mTEC revealed differentiating gene co-expression patterns that are maintained not only among individual mice of the same genetic background but also in different mouse strains, and that are also observed under different experimental designs (selected vs unselected) and batches. In aggregate, these results strongly suggest that an ordered process regulates PGE in mTEC.

Cell-based clustering of our data supports previous observations that the mTEC compartment is heterogeneous(Bornstein *et al*, 2018; Miragaia *et al*, 2018), a notion supported by studies tracking TEC differentiation during organogenesis and regeneration(Gäbler *et al*, 2007; Yano *et al*, 2008; Wang *et al*, 2012; Metzger *et al*, 2013; Nishikawa *et al*, 2014; Ohigashi *et al*, 2013; Mayer *et al*, 2016; Ohigashi *et al*, 2015). These suggest that mTEC derive during embryogenesis from progenitors expressing claudins-3 and −4 and SSEA-1(Hamazaki *et al*, 2007; Sekai *et al*, 2014), and postnatally from a population localised at the corticomedullary junction expressing podoplanin and CCL21(Onder *et al*, 2015; Michel *et al*, 2017; Mayer *et al*, 2016). Immature mTEC that are AIRE-CD80-CD86-MHCII^lo^, transition to an AIRE+CD80+CD86+MHCII^hi^ state that represents the most functionally mature mTEC from the perspective of PGE(Gäbler *et al*, 2007). Finally, mTEC lose expression of AIRE, MHCII and costimulatory molecules and enter a terminally differentiated state (Yano *et al*, 2008; Wang *et al*, 2012; Metzger *et al*, 2013; Nishikawa *et al*, 2014).

In keeping with the above model of mTEC maturation, two clusters (clusters 1 and 2) were identified that display progenitor-like characteristics because they express *Ccl21a* and *Pdpn* but lack *Aire, Cd80* and *Cd86* and are positive for transcripts encoding p63 (*Trp63*; Figure S5), a TEC lineage-specific determinant of proliferative capacity, *Itga6* (Cd49f; Figure S5), an integrin α-chain essential for TEC adhesion(Golbert *et al*, 2013), and *Sca-1* (Ly6a; Figure S5), a marker identifying a TEC progenitor status(Ulyanchenko *et al*, 2016; Wong *et al*, 2014). Custer 1 also expresses Eotaxin-1 (*Ccl11*; Figure S5), a chemokine known to attract eosinophils via the chemokine receptor Ccr3 suggesting a possible interaction between these mTEC and thymic resident eosinophils(Matthews *et al*, 1998; Throsby *et al*, 2000; Kim *et al*, 2010; Garcia-Zepeda *et al*, 1996). Pre-AIRE immature mTEC (cluster 2) transit to an *Aire+Cd80+Cd86+* (clusters 5 and 6) phenotype through an actively cycling stage (cluster 3), and then to a post-AIRE stage (clusters 7 and 8) with low *Aire, Cd80*, and *Cd86* and high *Krt10, Ivl* and *Spink5* expression.

Single-cell analyses recently defined distinct mTEC subpopulations. Bornstein *et al*.(Bornstein *et al*, 2018) identified four classes of mTEC (labelled mTEC I-IV) that largely agree with our data (Figure S3f). Clusters 1 and 2 relate to the immature pre-AIRE mTEC I subpopulation, clusters 3 - 6 to the mature AIRE+ mTEC II, clusters 7 and 8 to mTEC III, and cluster 10 to the tuft-like mTEC IV (Figure S3f)(Bornstein *et al*, 2018).

We also identified six novel mTEC clusters. Cluster 14 constitutes the largest newly described subpopulation (436 cells), enriched for mTEC expressing GP2 and defined by high expression of chemokine ligands *Ccl6* (Figure S5), *Ccl9* and *Ccl20*, as well as the chemokine receptor *Ccr5*. This result suggests that our selection of rare mTEC subtypes, using FACS enrichment, has revealed otherwise hidden subpopulations and thus that analyses of mTEC expressing other TRAs would likely uncover additional satellite clusters, each with a distinct transcriptome. Our second largest novel cluster is cluster 9 (227 cells), which expressed the markers *Ceacam10, Cd177* and *Ckm*. Cluster 9 shares transcriptome similarities not only with the terminally differentiated mTEC of cluster 8 but also with tuft-like mTEC of cluster 10. A third novel cluster, cluster 13 contains 117 mTEC and highly expresses several genes associated with cilium assembly (*Spag16, Wdr34* and *Bbs7*). The remaining novel clusters (clusters 11, 12 and 15) are defined by few cells (0.3 - 1% of the collected mTEC) and remain largely uncharacterised.

Low cell number has been a limitation of previous single-cell studies and is expected to have limited the detection of gene co-expression groups because of the low frequencies at which individual TRGs are expressed in single mTEC(Sansom *et al*, 2014; Meredith *et al*, 2015; Brennecke *et al*, 2015). In contrast to previous studies, which were either unable to detect co-expression(Sansom *et al*, 2014; Derbinski *et al*, 2008; Villaseñor *et al*, 2008), or were able only to detect a handful of co-expression groups(Meredith *et al*, 2015; Brennecke *et al*, 2015; Pinto *et al*, 2013), we were able to detect 50 separate gene co-expression modules, the majority of which comprised TRG co-expression sets (48 out of 50; Figure 5a; Figure 6a, b), suggesting that PGE in single mTEC follows an ordered process. Nevertheless, approximately one-third of the genes expressed in multiple cells could not be assigned to a co-expression group. Although this could reflect biological noise, it is possible that, with the inclusion of more single cells, these genes might also become assignable to specific gene modules.

A previous study found that co-expression patterns differed between individual mice and suggested that gene co-expression networks were established stochastically in each mouse(Meredith *et al*, 2015). This study investigated only 200 mTEC isolated from two pairs of WT or *Aire* knockout mice, raising a concern that it was significantly underpowered. By contrast, the gene modules identified in our study were robust as evidenced by their reproducibility across random samples of the data set (Figure 5b), as well as between individual mice independent of their strain (Figure 5c). Such reproducible TRG co-expression is likely to reflect an inherent cellular property that introduces bias in which TRGs are expressed together within a cell. That being said, we were unable to explain membership to co-expression groups by any of the features we examined, including gene category, chromosome localisation, genomic distance, tissue specificity, autoimmune disease association, and inclusion in a specific biological pathway. The largely random distribution of genes in modules across the genome is in keeping with observations from previous studies(Meredith *et al*, 2015; Pinto *et al*, 2013; Miragaia *et al*, 2018). Our results are also consistent with previously published studies that concluded that PGE patterns in mTEC are dictated neither by expression patterns seen in differentiated peripheral cell types nor by co-regulation by specific transcription factors(Meredith *et al*, 2015; Villaseñor *et al*, 2008; Pinto *et al*, 2013). Whilst we were unable to explain membership to co-expression groups this does not preclude the existence of an underlying mechanism that explains why certain TRGs are co-expressed in single mTEC particularly since inter-individual reproducibility was robust.

In the introduction we presented four molecular processes that could account for the heterogeneity of PGE in single mTEC. Our results provided evidence against TRG expression being entirely stochastic (Type 1) and also show that co-expression patterns in mTEC are not driven by the same cellular processes as in peripheral tissues (Type 3) or by contiguous colocation of co-expressed genes on the same chromosome (Type 4a). By contrast, our data provided evidence that different maturational stages or classes of mTEC activate TRG expression differentially (Type 2) and our data do not exclude the possibility that TRGs are physically co-located on chromatin by virtue of chromatin looping (Type 4b), as has beensuggested(Bansal *et al*, 2017; Pinto *et al*, 2013).

The biologically indeterminate yet ordered process for TRG co-expression described here suggests a system in which antigens are presented in a randomly dispersed manner across the medulla under a program that repeatedly generates mTEC with co-expressed genes that are randomly sampled with respect to disease-relevant antigens, pathways, tissues, and chromosomes. Furthermore, spatial location of GP2+ mTEC is randomly dispersed across the thymic medulla. This random spatial pattern of TRA presentation would provide a developing thymocyte travelling through a medullary island with the highest likelihood of encountering an mTEC expressing a given TRA against which its antigen receptor could be tested and implies that thymocytes would only need to traverse a limited volume within the thymic medulla in order to be tested against a diverse range of TRAs.

We conclude that reproducible order is evident among the genes that single mTEC express, yet the selection of these genes is indeterminate with respect to biological processes. This degree of randomness may ensure that single TEC express a wide range of TRAs, covering diverse peripheral tissues and auto-antigens, and that expression of a given TRA is spatially dispersed throughout the thymic medulla. In this way, a single thymocyte travelling through the thymic medulla may be given the greatest opportunity of encountering any given self-antigen for the purposes of central tolerance induction.

## Materials and Methods

### Mice

Female C57BL/6, BALB/c, and C57BL/6 x BALB/c F1 mice were obtained from Charles River Laboratories (Margate, Kent, UK) and rested for at least 1 week before analysis at 4-5 weeks of age. Mice were housed under specific pathogen-free conditions and according to institutional and UK Home Office regulations.

### Isolation of thymic epithelial cells and preparation for flow cytometry

TEC were isolated via enzymatic digestion of thymic lobes using Liberase (Roche) and DNaseI (Roche). Cells were counted and stained with anti-CD45 microbeads (Miltenyi Biotec) for 15 minutes at room temperature, before negative selection using the AutoMACS (Miltenyi Biotec) system in order to enrich for TEC. Samples were then stained for cell surface markers for 20 minutes at 4°C. For intracellular staining the Foxp3 Transcription Factor Staining Buffer Kit (eBioscience) was used according to manufacturer’s instructions. Combinations of UEA-1 lectin (Vector Laboratories) labelled in-house with Cy5 and the following antibodies were used to stain the cells: CD45::AF700 (30-F11, BioLegend), EpCAM::PerCPCy5.5 (G8.8, BioLegend), Ly51::PE (6C3, BioLegend), CD80::PECy5 (16-10A1, BioLegend), CD86::PECy7 (GL-1, BioLegend), MHCII::FITC (M5/114.15.2, BioLegend), MHCII::BV421 (M5/114.15.2, BioLegend), GP2::AF488 (2F11-C3, MBL), Rat IgG2a k::AF488 isotype control (eBR2a, eBioscience), TSPAN8::APC (657909, R&D Systems), Rat IgG2a k::APC isotype control (RTK4530, BioLegend), Desmoglein-3 (DSG3) unlabelled primary antibody (MBL) followed by secondary staining with goat anti-mouse IgG::APC-Cy7 (Abcam), AIRE::AF488 (5H12, eBioscience), AIRE::AF647 (5H12, eBioscience). For assessment of cell viability DAPI or the LIVE/DEAD Fixable Aqua Dead Cell Stain Kit were used (ThermoFisher Scientific). After staining cells were acquired and sorted using a FACS Aria III (BD Biosciences) and analysed using FlowJo v10 and GraphPad Prism 7; statistical analyses were performed using a t-test with Bonferroni correction for multiple comparisons where appropriate; differences were considered significant if the adjusted p-value was ≤ 0.05.

### RNA extraction, *reverse transcription and quantitative PCR (RT-qPCR)*

RNA was extracted using the RNeasy micro kit (Qiagen) and reverse transcribed using the SensiFAST cDNA synthesis kit (Bioline), according to manufacturer’s instructions. qPCR was then performed using the SensiFAST SYBR Hi-Rox kit (Bioline) and a StepOnePlus real-time PCR instrument (Applied Biosystems), using the following primer pairs (Sigma): *bActin* For 5’GTTCCGATGCCCTGAGGCTC3’, *bActin* Rev 5’CGGATGTCAACGTCACACTTCAT3’, *Gp2* For 5’CAAGAACAGATGCCCAAACCAA3’, *Gp2* Rev 5’AATGGCTGGTCTACTACTGCG3’, *Tspan8* For 5’TTCAGTCGGAGTTCAAGTGCT3’, *Tspan8* Rev 5’AACGGCCAGTCCAAAAGCAA3’. Data were analysed using GraphPad Prism 7; statistical analyses were performed using the t-test with Bonferroni correction for multiple comparisons; differences were considered significant if the adjusted p-value was ≤ 0.05.

### Single-cell RNA-sequencing

Cells were FACS sorted into 1.5ml DNase/RNase free Eppendorf tubes pre-coated with BSA and containing 150µl of plain RPMI-1640. Library preparation was carried out on fresh cells directly after FACS sorting using the Chromium Single Cell 3’ V1 kit or V2 kit (10X Genomics). The resulting libraries were sequenced on a HiSeq 2500 (Illumina) in High Output mode (paired-end asymmetric 100bp for read) or a HiSeq4000 (Illumina; paired-end 2×75bp).

### Analysis of single-cell RNA-sequencing results

Libraries were analyzed using the Cell Ranger pipeline (10X Genomics) resulting in a Gene-by-Barcode matrix of counts for each biological sample. The secondary analysis was performed using the simpleSingleCell workflow(Lun *et al*, 2016) (Bioconductor). Briefly, each dataset was filtered to remove low-quality libraries. These were defined as cells with a low number of reads (one median absolute deviation (MAD) lower than the median) or features (one MAD lower than the median), or a higher than expected percentage of reads from mitochondrial genes (three MADs higher than the median). The resulting gene-by-cell matrices for each experiment were normalised and the MNNcorrect algorithm(Haghverdi *et al*, 2018) was used to combine the separate experiments into one corrected meta-experiment.

Graph-based clustering was used to assign the individual mTEC to clusters. First, a shared nearest-neighbour (SNN) graph was generated from the principal components of the normalised gene-by-cell expression matrix. Community structure within the resulting SNN graph was then analysed using the Louvain algorithm(Blondel *et al*, 2008). This resulted in 15 high-quality clusters of mTEC from the combined meta-experiment.

mTEC were pseudo temporally ordered using two different schemes, firstly a diffusion map(Haghverdi *et al*, 2015) was used to get a reduced dimensionality representation of the data. That representation was used to order the cells along an inferred trajectory. Next, we used RNAvelocity (La Manno *et al*, 2018) to estimate the time derivative of the gene expression state of our mTEC thus ordering the mTEC based on spliced and unspliced RNA data captured from each cell.

To enhance the signal from the sparsely expressed TRGs and to prevent widely expressed genes from masking the signal of more sparsely expressed genes, gene module clustering was performed using an adaptation of the TF-IDF(Manning *et al*, 2008) transform. Firstly, a gene-frequency-by-inverse cell-frequency matrix was computed from the normalised gene-by-cell matrix. As this was a co-clustering analysis, only genes that were detected in multiple cells were included in the analysis. The gene frequency portion of the transform was computed as the log2 of normalised expression or the gene-by-cell expression matrix. The inverse cell-frequency was computed as the weighted average of the inverse frequency of detection of each gene within each subpopulation. That is, for gene *X* in subset *Y*: if *X* is detected in 25% of *Y* then the inverse cell-frequency is 4. The five conditions (TSPAN8+/-, GP2+/- or Unselected) were weighted by their expected contribution to the total mTEC population (Figure 1a) and the resulting average inverse cell-frequency log10 transformed before the product of the gene-frequency matrix and inverse cell-frequency were computed. The gene-frequency-by-inverse cell-frequency matrix was further reduced to a gene-by-context matrix by using a t-distributed stochastic neighbour embedding (t-SNE)(Maaten & Hinton, 2008) to reduce the cosine distance of the first 50 eigenvectors of the gene-frequency-by-inverse cell-frequency matrix (acquired using singular value decomposition). This reduced dimensionality gene-by-context matrix was then clustered using HDBSCAN(McInnes & Healy, 2017) to spatially select clusters based on density in the reduced dimensionality representation. This has the benefit of identifying the sets of genes that are repeatedly observed together in the same context (subsets of cells), while simultaneously attenuating the signal from frequently expressed genes unless accompanied by a drastic change in expression level.

Co-expression modules were tested for robustness by repeating the gene module clustering on 100 random subsets of the meta-experiment. The gene modules identified from the random subsets were then compared to determine the overall similarity in the assignment of genes to modules. An adjusted mutual information (AMI) score was used to compare each set of assignments in a pairwise fashion. The clustering algorithm we used either assigns each gene to a module or declares it as noise (unassigned). To compare the clusterings the AMI was calculated using the unassigned/noise cells as a cluster.

As a secondary analysis to quantify gene co-expression, we calculated the pairwise co-expression frequency of all TRGs within individual mice. This frequency f(G^X^,G^Y^) was computed as the fraction of cells from a single mouse in which both gene X and gene Y were detected. These frequencies were compared across all mice and to the full meta-experiment (all cells) using a Pearson correlation.

We examined multiple features of co-expressed genes in order to determine whether a particular feature was able to explain the co-expression patterns that we observe. Firstly, we considered whether each module contains the expected proportion of TRGs or if some module was enriched for TRGs. In this analysis, a Monte Carlo simulation was used to generate the expected number of TRGs in each module and an empirical p-value was generated for each observed value. Next, we examined the location of genes within each module to determine if any modules prefer a particular chromosome. Accordingly, we computed the percentage of genes within a given module that are encoded on each chromosome. Finally, we used a Monte Carlo simulation to determine if this percentage was more extreme than expected by chance. On a more local scale, we next sought to determine if co-expressed genes were clustered closer than expected by chance (within chromosomes). To determine this, we computed the full pairwise distance matrix between all pairs of co-expressed genes. We compared the pairwise distance distribution of all genes to only pairs of AIRE-regulated genes or to pairs of genes from the same gene expression module and found very little difference between those distributions and the full distribution for all genes.

### Immunohistochemistry and Confocal Microscopy

Freshly isolated thymic lobes were frozen in OCT compound (Tissue-Tek) and cryosectioned at a thickness of 8µm. For immunofluorescence staining, tissue sections were fixed in 10% neutral buffered formalin (Sigma) for 20 minutes at room temperature and then permeabilised in PBS 0.3% Triton X-100 (Sigma) for 10 minutes. This was followed by incubation for 1 hour at room temperature in blocking buffer consisting of 2% goat serum (Sigma) in PBS 0.1% Triton X-100. The slides were then stained with a rabbit primary anti-cytokeratin 5 (KRT5) antibody (Covance/BioLegend) diluted 1:500 in blocking buffer for 1 hour at 37°C, after which three 5 minute PBS washing steps were performed. Secondary antibody staining with goat anti-rabbit::AF555 diluted 1:500 in PBS 0.1% Triton X-100 was carried out for 30 minutes at 37°C, followed by three 5 minute PBS washing steps. A third staining step with a 1:200 dilution of an anti-GP2 antibody directly conjugated to AF488 (MBL, 2F11-C3) or an isotype control (Rat IgG2a k::AF488 isotype control, eBR2a, eBioscience) was subsequently performed for 1 hour at 37°C. After washing as above, the slides were stained using 500ng/ml DAPI (Sigma) diluted in methanol (VWR), washed once in PBS and mounted using ProLong Gold Antifade mounting medium (Life Technologies). Imaging was performed on an LSM 780 inverted confocal microscope (Ziess) and analysed using Fiji (Schindelin *et al*, 2012).

### Spatial analysis of GP2+ distribution

Four representative 1980×1980 pixel microscope images co-stained with DAPI, GP2 and KRT5 were used to identify GP2+ mTEC and the medullary region within thymic images. Each colour image was opened and processes using the EBImage package (R;(Pau *et al*, 2010)). KRT5 layers were processed with a low pass Gaussian filter then thresholded images were eroded and dilated to obtain a mask of the region covered by mTEC. A similar process was used to identify GP2+ mTEC on the GP2+ layer, an additional stage of watershed processing completed the labels of individual GP2+ mTEC. The moment of each GP2+ feature was computed and validated by visual inspection and these positions were used in the spatial analysis. To determine if the GP2+ mTEC were clustered or randomly dispersed within the medullary region the nearest neighbour distance distribution function (G(r): spatstat package R) was calculated for the point pattern derived from the moments of the GP2+ mTEC within the space classified as medulla by the KRT5+ mask generated above.

## Data availability

These data are available through ArrayExpress under the ID TBD.

## Acknowledgements

This work was supported by the Wellcome Trust [109032/Z/15/Z], [105045/Z/14/2] and the MRC [MC_UU_00007/15 to CPP].

## Contributions

FD, SM, JB, GH and CPP designed the experiments. FD and SM conducted the experiments; LC helped with 10X library preparation. FD and JB analysed the data and produced the figures. FD, JB, GH and CPP contributed to writing the manuscript.

## References

Abramson J & Anderson G (2017) Thymic Epithelial Cells. Annu. Rev. Immunol. 35: 85–118

Agaësse G, Barbollat-Boutrand L, El Kharbili M, Berthier-Vergnes O & Masse I (2017) p53 targets TSPAN8 to prevent invasion in melanoma cells. Oncogenesis 6: e309

Bansal K, Yoshida H, Benoist C & Mathis D (2017) The transcriptional regulator Aire binds to and activates super-enhancers. Nat. Immunol. 18: 263–273

Barthlott T, Keller MP, Krenger W & Holländer GA (2006) A short primer on early molecular and cellular events in thymus organogenesis and replacement. Swiss Med. Wkly 136: 365–369

Bitoun E, Micheloni A, Lamant L, Bonnart C, Tartaglia-Polcini A, Cobbold C, Al Saati T, Mariotti F, Mazereeuw-Hautier J, Boralevi F, Hohl D, Harper J, Bodemer C, D’Alessio M & Hovnanian A (2003) LEKTI proteolytic processing in human primary keratinocytes, tissue distribution and defective expression in Netherton syndrome. Hum. Mol. Genet. 12: 2417–2430

Blondel VD, Guillaume J-L, Lambiotte R & Lefebvre E (2008) Fast unfolding of communities in large networks. arXiv [physics.soc-ph] Available at: https://arxiv.org/abs/0803.0476

Bornstein C, Nevo S, Giladi A, Kadouri N, Pouzolles M, Gerbe F, David E, Machado A, Chuprin A, Tóth B, Goldberg O, Itzkovitz S, Taylor N, Jay P, Zimmermann VS, Abramson J & Amit I (2018) Single-cell mapping of the thymic stroma identifies IL-25-producing tuft epithelial cells. Nature 559: 622–626

Brennecke P, Reyes A, Pinto S, Rattay K, Nguyen M, Küchler R, Huber W, Kyewski B & Steinmetz LM (2015) Single-cell transcriptome analysis reveals coordinated ectopic gene-expression patterns in medullary thymic epithelial cells. Nat. Immunol. 16: 933–941

Cogger KF, Sinha A, Sarangi F, McGaugh EC, Saunders D, Dorrell C, Mejia-Guerrero S, Aghazadeh Y, Rourke JL, Screaton RA, Grompe M, Streeter PR, Powers AC, Brissova M, Kislinger T & Nostro MC (2017) Glycoprotein 2 is a specific cell surface marker of human pancreatic progenitors. Nat. Commun. 8: 331

Derbinski J, Gäbler J, Brors B, Tierling S, Jonnakuty S, Hergenhahn M, Peltonen L, Walter J & Kyewski B (2005) Promiscuous gene expression in thymic epithelial cells is regulated at multiple levels. J. Exp. Med. 202: 33–45

Derbinski J, Pinto S, Rösch S, Hexel K & Kyewski B (2008) Promiscuous gene expression patterns in single medullary thymic epithelial cells argue for a stochastic mechanism. Proc. Natl. Acad. Sci. U. S. A. 105: 657–662

Derbinski J, Schulte A, Kyewski B & Klein L (2001) Promiscuous gene expression in medullary thymic epithelial cells mirrors the peripheral self. Nat. Immunol. 2: 1032–1039

Gäbler J, Arnold J & Kyewski B (2007) Promiscuous gene expression and the developmental dynamics of medullary thymic epithelial cells. Eur. J. Immunol. 37: 3363–3372

Galliano MF, Roccasecca RM, Descargues P, Micheloni A, Levy E, Zambruno G, D’Alessio M & Hovnanian A (2005) Characterization and expression analysis of the Spink5 gene, the mouse ortholog of the defective gene in Netherton syndrome. Genomics 85: 483–492

Garcia-Zepeda EA, Rothenberg ME, Ownbey RT, Celestin J, Leder P & Luster AD (1996) Human eotaxin is a specific chemoattractant for eosinophil cells and provides a new mechanism to explain tissue eosinophilia. Nat. Med. 2: 449–456

Golbert DCF, Correa-de-Santana E, Ribeiro-Alves M, de Vasconcelos ATR & Savino W (2013) ITGA6 gene silencing by RNA interference modulates the expression of a large number of cell migration-related genes in human thymic epithelial cells. BMC Genomics 14 Suppl 6: S3

Haghverdi L, Buettner F & Theis FJ (2015) Diffusion maps for high-dimensional single-cell analysis of differentiation data. Bioinformatics 31: 2989–2998

Haghverdi L, Lun ATL, Morgan MD & Marioni JC (2018) Batch effects in single-cell RNA-sequencing data are corrected by matching mutual nearest neighbors. Nat. Biotechnol. 36: 421–427

Hamazaki Y, Fujita H, Kobayashi T, Choi Y, Scott HS, Matsumoto M & Minato N (2007) Medullary thymic epithelial cells expressing Aire represent a unique lineage derived from cells expressing claudin. Nat. Immunol. 8: 304–311

Hogquist KA & Jameson SC (2014) The self-obsession of T cells: how TCR signaling thresholds affect fate ‘decisions’ and effector function. Nat. Immunol. 15: 815–823

Kim H-J, Alonzo ES, Dorothee G, Pollard JW & Sant’Angelo DB (2010) Selective depletion of eosinophils or neutrophils in mice impacts the efficiency of apoptotic cell clearance in the thymus. PLoS One 5: e11439

Klein L, Hinterberger M, Wirnsberger G & Kyewski B (2009) Antigen presentation in the thymus for positive selection and central tolerance induction. Nat. Rev. Immunol. 9: 833–844

Klein L, Kyewski B, Allen PM & Hogquist KA (2014) Positive and negative selection of the T cell repertoire: what thymocytes see (and don’t see). Nat. Rev. Immunol. 14: 377–391

Kyewski B & Klein L (2006) A central role for central tolerance. Annu. Rev. Immunol. 24: 571–606

La Manno G, Soldatov R, Zeisel A, Braun E, Hochgerner H, Petukhov V, Lidschreiber K, Kastriti ME, Lönnerberg P, Furlan A, Fan J, Borm LE, Liu Z, van Bruggen D, Guo J, He X, Barker R, Sundström E, Castelo-Branco G, Cramer P, et al (2018) RNA velocity of single cells. Nature 560: 494–498

Lun ATL, McCarthy DJ & Marioni JC (2016) A step-by-step workflow for low-level analysis of single-cell RNA-seq data with Bioconductor. F1000Res. 5: 2122

Maaten L van der & Hinton G (2008) Visualizing Data using t-SNE. J. Mach. Learn. Res. 9: 2579–2605

Manning CD, Raghavan P & Schütze H (2008) Introduction to Information Retrieval Cambridge University Press

Matthews AN, Friend DS, Zimmermann N, Sarafi MN, Luster AD, Pearlman E, Wert SE & Rothenberg ME (1998) Eotaxin is required for the baseline level of tissue eosinophils. Proc. Natl. Acad. Sci. U. S. A. 95: 6273–6278

Mayer CE, Žuklys S, Zhanybekova S, Ohigashi I, Teh H-Y, Sansom SN, Shikama-Dorn N, Hafen K, Macaulay IC, Deadman ME, Ponting CP, Takahama Y & Holländer GA (2016) Dynamic spatio-temporal contribution of single β5t+ cortical epithelial precursors to the thymus medulla. Eur. J. Immunol. 46: 846–856

McInnes L & Healy J (2017) Accelerated Hierarchical Density Based Clustering. In 2017 IEEE International Conference on Data Mining Workshops (ICDMW) pp 33–42.

Meredith M, Zemmour D, Mathis D & Benoist C (2015) Aire controls gene expression in the thymic epithelium with ordered stochasticity. Nat. Immunol. 16: 942–949

Metzger TC, Khan IS, Gardner JM, Mouchess ML, Johannes KP, Krawisz AK, Skrzypczynska KM & Anderson MS (2013) Lineage Tracing and Cell Ablation Identify a Post-Aire-Expressing Thymic Epithelial Cell Population. Cell Rep. 5: 166–179

Michel C, Miller CN, Küchler R, Brors B, Anderson MS, Kyewski B & Pinto S (2017) Revisiting the Road Map of Medullary Thymic Epithelial Cell Differentiation. J. Immunol. Available at: http://dx.doi.org/10.4049/jimmunol.1700203

Miller CN, Proekt I, von Moltke J, Wells KL, Rajpurkar AR, Wang H, Rattay K, Khan IS, Metzger TC, Pollack JL, Fries AC, Lwin WW, Wigton EJ, Parent AV, Kyewski B, Erle DJ, Hogquist KA, Steinmetz LM, Locksley RM & Anderson MS (2018) Thymic tuft cells promote an IL-4-enriched medulla and shape thymocyte development. Nature 559: 627–631

Miragaia RJ, Zhang X, Gomes T, Svensson V, Ilicic T, Henriksson J, Kar G & Lönnberg T (2018) Single-cell RNA-sequencing resolves self-antigen expression during mTEC development. Sci. Rep. 8: 685

Nishikawa Y, Hirota F, Yano M, Kitajima H, Miyazaki J-I, Kawamoto H, Mouri Y & Matsumoto M (2010) Biphasic Aire expression in early embryos and in medullary thymic epithelial cells before end-stage terminal differentiation. J. Exp. Med. 207: 963–971

Nishikawa Y, Nishijima H, Matsumoto M, Morimoto J, Hirota F, Takahashi S, Luche H, Fehling HJ, Mouri Y & Matsumoto M (2014) Temporal lineage tracing of Aire-expressing cells reveals a requirement for Aire in their maturation program. J. Immunol. 192: 2585–2592

Ohigashi I, Zuklys S, Sakata M, Mayer CE, Hamazaki Y, Minato N, Hollander GA & Takahama Y (2015) Adult Thymic Medullary Epithelium Is Maintained and Regenerated by Lineage-Restricted Cells Rather Than Bipotent Progenitors. Cell Rep. 13: 1432–1443

Ohigashi I, Zuklys S, Sakata M, Mayer CE, Zhanybekova S, Murata S, Tanaka K, Holländer GA & Takahama Y (2013) Aire-expressing thymic medullary epithelial cells originate from β5t-expressing progenitor cells. Proc. Natl. Acad. Sci. U. S. A. 110: 9885–9890

Ohno H & Hase K (2010) Glycoprotein 2 (GP2): grabbing the FimH bacteria into M cells for mucosal immunity. Gut Microbes 1: 407–410

Onder L, Nindl V, Scandella E, Chai Q, Cheng H-W, Caviezel-Firner S, Novkovic M, Bomze D, Maier R, Mair F, Ledermann B, Becher B, Waisman A & Ludewig B (2015) Alternative NF-κB signaling regulates mTEC differentiation from podoplanin-expressing precursors in the cortico-medullary junction. Eur. J. Immunol. 45: 2218–2231

Pau G, Fuchs F, Sklyar O, Boutros M & Huber W (2010) EBImage-an R package for image processing with applications to cellular phenotypes. Bioinformatics 26: 979–981

Pinto S, Michel C, Schmidt-Glenewinkel H, Harder N, Rohr K, Wild S, Brors B & Kyewski B (2013) Overlapping gene coexpression patterns in human medullary thymic epithelial cells generate self-antigen diversity. Proc. Natl. Acad. Sci. U. S. A. 110: E3497–505

Rattay K, Meyer HV, Herrmann C, Brors B & Kyewski B (2016) Evolutionary conserved gene co-expression drives generation of self-antigen diversity in medullary thymic epithelial cells. J. Autoimmun. 67: 65–75

Reid L, Meyrick B, Antony VB, Chang L-Y, Crapo JD & Reynolds HY (2005) The mysterious pulmonary brush cell: a cell in search of a function. Am. J. Respir. Crit. Care Med. 172: 136–139

Rodewald H-R (2008) Thymus organogenesis. Annu. Rev. Immunol. 26: 355–388

Sansom SN, Shikama-Dorn N, Zhanybekova S, Nusspaumer G, Macaulay IC, Deadman ME, Heger A, Ponting CP & Holländer GA (2014) Population and single-cell genomics reveal the Aire dependency, relief from Polycomb silencing, and distribution of self-antigen expression in thymic epithelia. Genome Res. 24: 1918–1931

Schindelin J, Arganda-Carreras I, Frise E, Kaynig V, Longair M, Pietzsch T, Preibisch S, Rueden C, Saalfeld S, Schmid B, Tinevez J-Y, White DJ, Hartenstein V, Eliceiri K, Tomancak P & Cardona A (2012) Fiji: an open-source platform for biological-image analysis. Nat. Methods 9: 676–682

Scialdone A, Natarajan KN, Saraiva LR, Proserpio V, Teichmann SA, Stegle O, Marioni JC & Buettner F (2015) Computational assignment of cell-cycle stage from single-cell transcriptome data. Methods 85: 54–61

Sekai M, Hamazaki Y & Minato N (2014) Medullary thymic epithelial stem cells maintain a functional thymus to ensure lifelong central T cell tolerance. Immunity 41: 753–761

Sun L, Luo H, Li H & Zhao Y (2013) Thymic epithelial cell development and differentiation: cellular and molecular regulation. Protein Cell 4: 342–355

Takahama Y (2006) Journey through the thymus: stromal guides for T-cell development and selection. Nat. Rev. Immunol. 6: 127–135

Throsby M, Herbelin A, Pléau JM & Dardenne M (2000) CD11c+ eosinophils in the murine thymus: developmental regulation and recruitment upon MHC class I-restricted thymocyte deletion. J. Immunol. 165: 1965–1975

Todd R & Wong DT (1999) Oncogenes. Anticancer Res. 19: 4729–4746

Tornai T, Tornai D, Sipeki N, Tornai I, Alsulaimani R, Fechner K, Roggenbuck D, Norman GL, Veres G, Par G, Par A, Szalay F, Lakatos PL, Antal-Szalmas P & Papp M (2018) Loss of tolerance to gut immunity protein, glycoprotein 2 (GP2) is associated with progressive disease course in primary sclerosing cholangitis. Sci. Rep. 8: 399

Ulyanchenko S, O’Neill KE, Medley T, Farley AM, Vaidya HJ, Cook AM, Blair NF & Blackburn CC (2016) Identification of a Bipotent Epithelial Progenitor Population in the Adult Thymus. Cell Rep. 14: 2819–2832

Vaidya HJ, Briones Leon A & Blackburn CC (2016) FOXN1 in thymus organogenesis and development. Eur. J. Immunol. 46: 1826–1837

Villaseñor J, Besse W, Benoist C & Mathis D (2008) Ectopic expression of peripheral-tissue antigens in the thymic epithelium: probabilistic, monoallelic, misinitiated. Proc. Natl. Acad. Sci. U. S. A. 105: 15854–15859

Wada N, Nishifuji K, Yamada T, Kudoh J, Shimizu N, Matsumoto M, Peltonen L, Nagafuchi S & Amagai M (2011) Aire-dependent thymic expression of desmoglein 3, the autoantigen in pemphigus vulgaris, and its role in T-cell tolerance. J. Invest. Dermatol. 131: 410–417

Wang X, Laan M, Bichele R, Kisand K, Scott HS & Peterson P (2012) Post-Aire maturation of thymic medullary epithelial cells involves selective expression of keratinocyte-specific autoantigens. Front. Immunol. 3: 19

Werner L, Sturm A, Roggenbuck D, Yahav L, Zion T, Meirowithz E, Ofer A, Guzner-Gur H, Tulchinsky H & Dotan I (2013) Antibodies against glycoprotein 2 are novel markers of intestinal inflammation in patients with an ileal pouch. J. Crohns. Colitis 7: e522–32

Wong K, Lister NL, Barsanti M, Lim JMC, Hammett MV, Khong DM, Siatskas C, Gray DHD, Boyd RL & Chidgey AP (2014) Multilineage potential and self-renewal define an epithelial progenitor cell population in the adult thymus. Cell Rep. 8: 1198–1209

Yamashita J, Ohmoto M, Yamaguchi T, Matsumoto I & Hirota J (2017) Skn-1a/Pou2f3 functions as a master regulator to generate Trpm5-expressing chemosensory cells in mice. PLoS One 12: e0189340

Yano M, Kuroda N, Han H, Meguro-Horike M, Nishikawa Y, Kiyonari H, Maemura K, Yanagawa Y, Obata K, Takahashi S, Ikawa T, Satoh R, Kawamoto H, Mouri Y & Matsumoto M (2008) Aire controls the differentiation program of thymic epithelial cells in the medulla for the establishment of self-tolerance. J. Exp. Med. 205: 2827–2838

Zhao K, Erb U, Hackert T, Zöller M & Yue S (2018) Distorted leukocyte migration, angiogenesis, wound repair and metastasis in Tspan8 and Tspan8/CD151 double knockout mice indicate complementary activities of Tspan8 and CD51. Biochim. Biophys. Acta Mol. Cell Res. 1865: 379–391

Zhu Y, Ailane N, Sala-Valdés M, Haghighi-Rad F, Billard M, Nguyen V, Saffroy R, Lemoine A, Rubinstein E, Boucheix C & Greco C (2017) Multi-factorial modulation of colorectal carcinoma cells motility-partial coordination by the tetraspanin Co-029/tspan8. Oncotarget 8: 27454–27470

